# Keratinocyte desmosomal cadherin Desmoglein 1 as a mediator and target of paracrine signaling in the melanoma niche

**DOI:** 10.1101/2022.12.01.518424

**Authors:** Hope E. Burks, Jenny L. Pokorny, Jennifer L. Koetsier, Quinn R. Roth-Carter, Christopher R. Arnette, Pedram Gerami, John T. Seykora, Jodi L. Johnson, Kathleen J. Green

## Abstract

Melanoma is an aggressive cancer typically arising from transformation of melanocytes residing in the basal layer of the epidermis, where they are in direct contact with surrounding keratinocytes. The role of keratinocytes in shaping the melanoma tumor microenvironment remains understudied. We previously showed that temporary loss of the keratinocyte-specific cadherin, Desmoglein 1 (Dsg1) controls paracrine signaling between normal melanocytes and keratinocytes to stimulate the protective tanning response. Here, we provide evidence that melanoma cells hijack this intercellular communication by secreting factors that keep Dsg1 expression low in surrounding keratinocytes, which in turn generate their own paracrine signals that enhance melanoma spread through CXCL1/CXCR2 signaling. Evidence suggests a model whereby paracrine signaling from melanoma cells increases levels of the transcriptional repressor *Snai2* (Slug), and consequently decreases the Dsg1 transcriptional activator *Grhl1* (Grainyhead like 1), a known promoter of epidermal differentiation and barrier function. Together, these data support the idea that paracrine crosstalk between melanoma cells and keratinocytes resulting in chronic keratinocyte Dsg1 reduction contributes to melanoma cell movement associated with tumor progression.

## Introduction

Cutaneous melanoma is a deadly skin cancer arising from pigmented melanocytes within the epidermis. Interactions between melanoma cells and their surrounding microenvironment play a critical role in tumor initiation and progression (Brandner and Haass, 2013; Villanueva and Herlyn, 2008; Wang et al., 2016). Discoveries surrounding immune cell interactions in solid tumors have led to significant innovation in clinical treatment strategies across multiple malignancies including melanoma (Kalaora et al., 2022). Similarly, the role of fibroblast-melanoma cell interactions has been integral to understanding melanoma spread within the dermis (Galbo et al., 2021). However, less is known about how communication between melanocytes and their more abundant keratinocyte neighbors promotes melanoma development or initial stages of melanoma movement within and away from the epidermal environment.

It is well known that melanocyte-keratinocyte communication is crucial for mediating protective responses to ultraviolet radiation (UV) (Upadhyay et al., 2021; Wang et al., 2016; Yardman-Frank and Fisher, 2021). UV elicits a cascade of signaling between the two cell types, resulting in the production of melanin by melanocytes, which is transferred to keratinocytes, where the pigment is positioned over the nucleus to protect DNA from UV-mediated damage. Changes in communication between pigmented cells and keratinocytes continuously evolve in the developing melanoma environment (Brandner and Haass, 2013). Melanoma cells can escape control by adjacent keratinocytes by losing their intercellular contacts, allowing uncontrolled proliferation and spread (Wang et al., 2016). Cell contact between radially spreading melanoma cells and differentiated keratinocytes can then trigger vertical growth through Notch signaling, promoting movement of melanoma cells into the dermis (Golan et al., 2015). Major contributors to cell contact-mediated communication between keratinocytes and melanocytes are members of the cadherin family of cell-cell adhesion molecules. For instance, loss of E-cadherin (Ecad) is associated with the loss of keratinocyte control over melanoma cell proliferation and invasion (Haass et al., 2005; Hsu et al., 2000). Further, increased P-cadherin stabilizes heterotypic adhesions between keratinocytes and melanoma cells to promote melanoma cell expansion and is associated with reduced survival in humans (Mescher et al., 2017).

Beyond cadherins’ functions in direct contact, a role for vertebrate-specific cadherins known as desmogleins in paracrine signaling has recently emerged. Desmogleins work with partner desmocollin proteins to make up the adhesive core of desmosomes, essential intercellular junctions that provide structural integrity to the skin and are critical for epidermal barrier function (Hegazy et al., 2022; Kowalczyk and Green, 2013). Desmoglein 1 (Dsg1) is a keratinocyte specific desmosomal cadherin found in complex stratified tissues of terrestrial organisms (Green et al., 2020). Work in our lab has shown that Dsg1 plays roles in cell signaling independent of its role as an adhesion molecule. Dsg1 promotes keratinocyte differentiation through suppression of EGFR-MAPK signaling by associating with Erbin, ultimately inhibiting Ras-Raf complex formation upstream of ERK1/2 to promote differentiation (Harmon et al., 2013). Dsg1 also controls cytokine signaling to dampen inflammation, and its loss in patients with bi-allelic mutations in Dsg1 results in the inflammatory condition severe dermatitis, multiple allergies and metabolic wasting (SAM) syndrome (Godsel et al., 2022; Polivka et al., 2018; Samuelov et al., 2013).

We recently showed that paracrine signaling controlled by keratinocyte Dsg1 also controls melanocyte behavior (Arnette et al., 2019). Dsg1, but not Ecad, is temporarily lost in a dose dependent manner following acute UV radiation (Johnson et al., 2014). Modeling this behavior, genetic interference resulting in acute loss of Dsg1 affects melanocyte pigmentation and dendricity in a manner reminiscent of the tanning response through secretion of a subset of cytokines also known to be generated in keratinocytes stimulated with UV (Arnette et al., 2019). While acute loss of Dsg1 seems to play a protective role in skin by activating the tanning response, long-term loss of keratinocyte Dsg1 promotes behaviors in melanocytes that mimic a malignant phenotype, such as pagetoid movement in 3D organotypic cultures (Arnette et al., 2019). These observations reveal a potential role for loss of keratinocyte Dsg1 in initiating metastatic spread in the primary melanoma niche, specifically in intra-epidermal movement, an understudied phenomenon in melanoma progression.

Here, through a combination of analyses of human melanoma tissues and *in vitro* mechanistic studies, we identify a bi-directional communication pathway between keratinocytes and melanoma cells. This communication depends on the melanoma cells’ ability to reduce Dsg1 expression. Consequent keratinocyte-dependent paracrine stimulation of a neural crest like signature in melanoma cells associates with increased melanoma cell migration. Our data support the idea that melanoma cells reduce Dsg1 by producing paracrine factors that increase activity of the transcription factor Slug and decrease levels of the Dsg1 transcription factor Grainyhead like 1 (Grhl1). The relevance of these changes is supported by analysis of human melanoma sections showing that persistent loss of Dsg1, but not Ecad, is associated with local melanoma spread, reduced Grhl1, and increased Slug. Based on these observations, we propose that melanomas hijack normal keratinocyte to melanocyte crosstalk to maintain a pro-tumor environment.

## Results

### Desmoglein 1 is reduced in melanoma adjacent keratinocytes in human epidermis

To address whether there is an association between Dsg1 loss and melanoma, benign and malignant melanocytic lesions were stained for Dsg1, Ecad in keratinocytes and Melan-A (Mel-A) to mark melanoma cells (Fig. 1A). Quantifying Dsg1 and Ecad border localization and staining intensity revealed that Dsg1 was significantly reduced in the keratinocytes directly adjacent to the Mel-A positive staining cells (melanocytes/melanoma cells) in melanoma samples compared with distant sites or control tissues (Fig. 1B, C). In contrast, keratinocytes adjacent to melanocytes in benign nevi did not show any alteration in Dsg1 levels. Ecad staining remained unchanged in all conditions, demonstrating that the loss of Dsg1 is not solely attributable to a general loss of cell adhesion proteins in melanoma adjacent keratinocytes (Fig. 1C).

**Figure 1:**
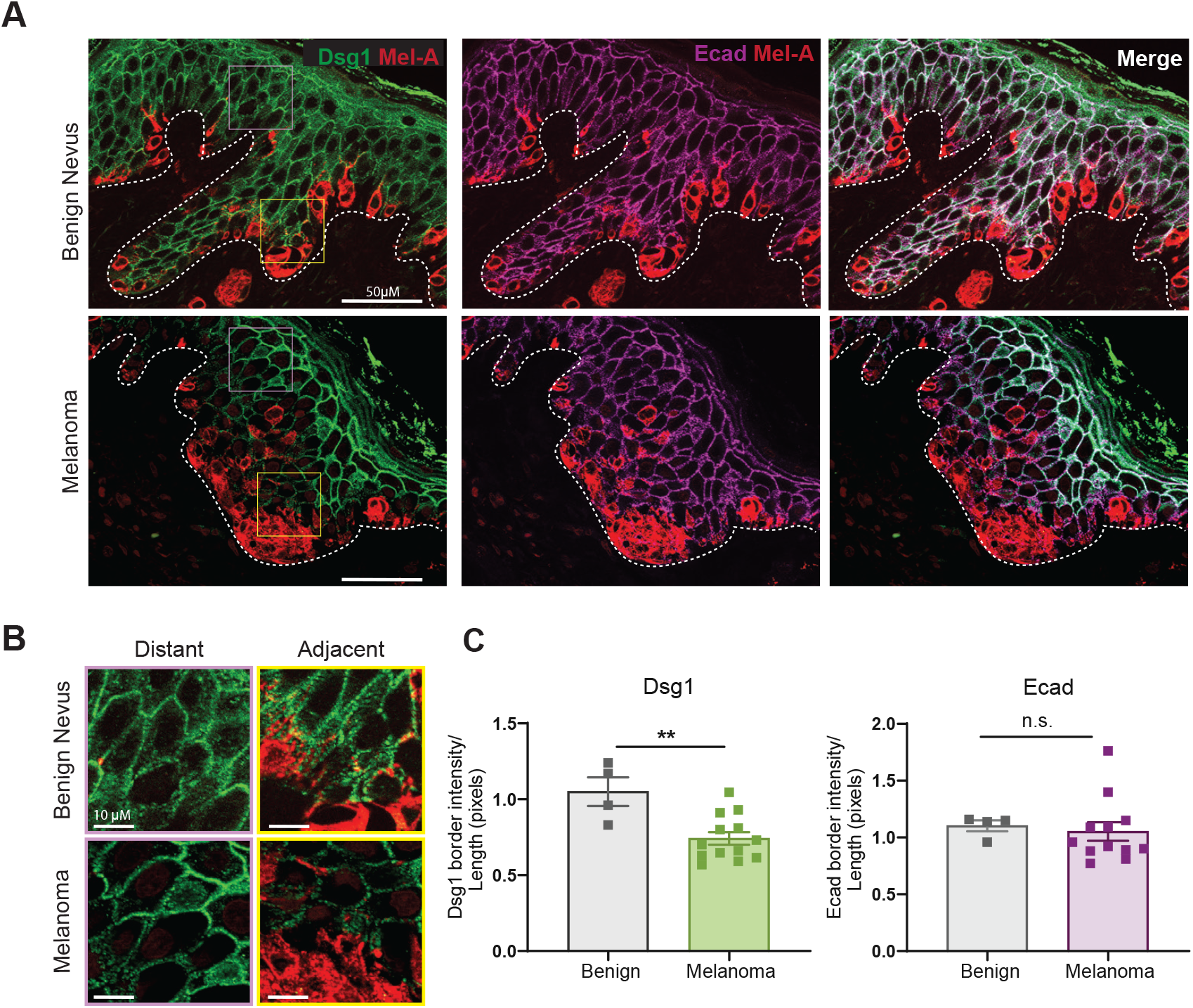
Desmoglein 1 is reduced in melanoma adjacent keratinocytes in human epidermis. A) Paraffin-embedded sections of benign nevi and primary stage melanoma were co-stained for desmoglein 1 (Dsg1) or Ecad and Mel-A to highlight cadherins at cell-cell interfaces of keratinocytes surrounding pigmented cells. Dashed line indicates basement membrane. B) Both distant (2-4 cells away) and proximal keratinocytes were examined for cadherin staining for quantification. C) Pixel intensities were determined for 4 benign and 12 melanoma samples and the ratio of distant to proximal intensities plotted. A significant decrease in Dsg1, but not Ecad, intensity proximal to the melanoma lesions was observed. Mean +/- SEM depicted. **p< 0.01. Student’s t-test.

### Melanoma cells downregulate keratinocyte Dsg1 through paracrine signaling

Comparisons of chronic sun exposed skin and non-sun exposed skin transcriptomes from the Genotype-Tissue Expression (GTEx) database showed no significant change in Dsg1 gene expression levels (Fig. S1). This, combined with our previous finding that Dsg1 levels recover over time after UV exposure (Johnson et al., 2014), suggests that sustained loss of Dsg1 is likely a disease specific phenomenon and not the result of accumulated sun damage. Furthermore, Dsg1 is maintained in keratinocytes distant from lesional melanoma cells. This led us to ask whether keratinocyte Dsg1 is regulated by the adjacent melanoma cells.

To address this question, we began with a unidirectional approach using conditioned media from a panel of melanoma cell lines varying in growth phase and mutation status. Differentiated keratinocytes were cultured in conditioned media from control primary melanocytes or melanoma cell lines, and mRNA or protein was collected and assessed by RT-PCR or immunoblot, respectively. The mRNA and protein levels of Dsg1, but not Ecad or Desmoglein 3 (Dsg3), another desmosomal cadherin, were decreased in keratinocytes cultured in melanoma but not melanocyte conditioned media (Fig. 2A, B).

**Figure 2:**
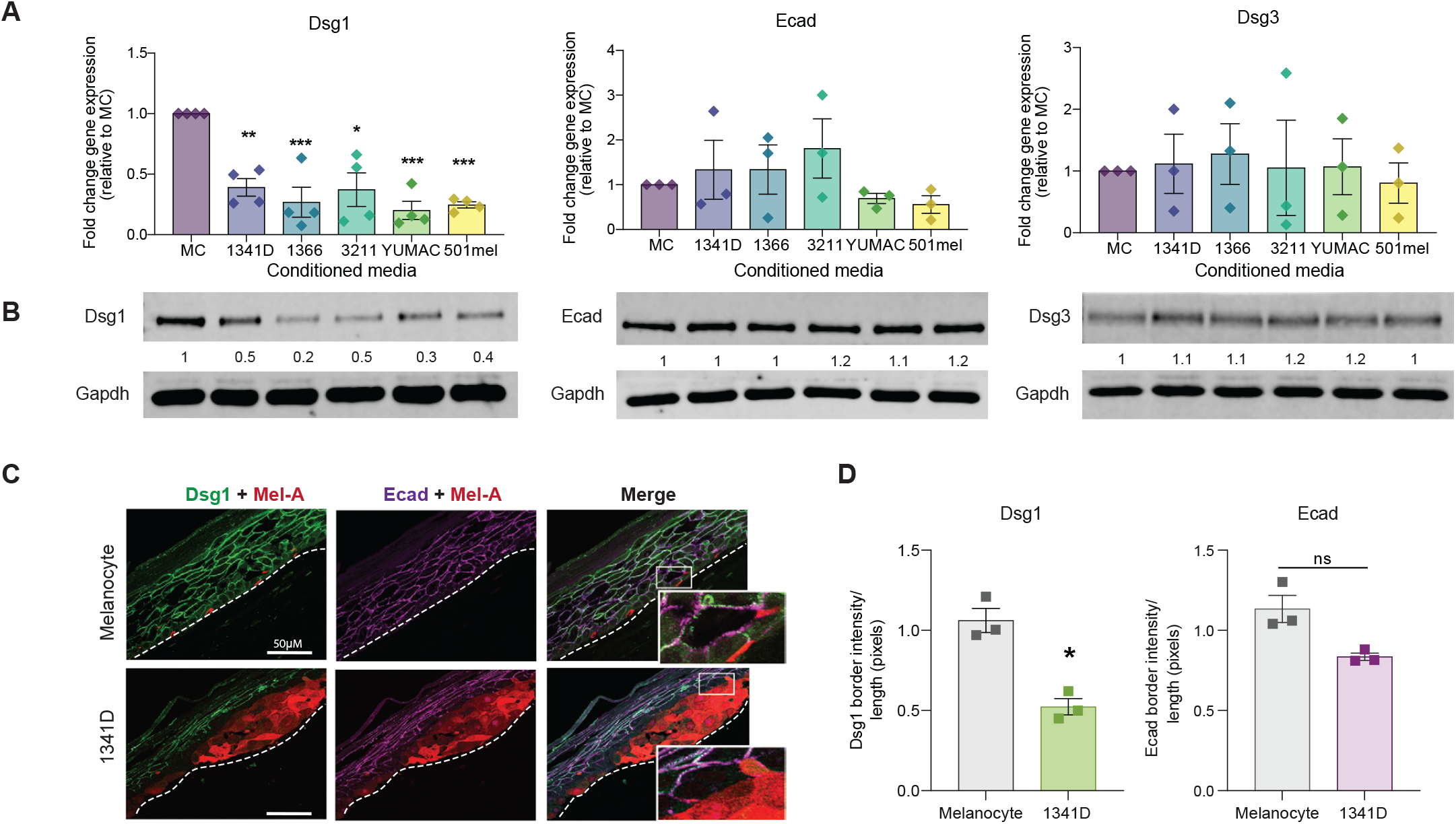
Melanoma cells downregulate keratinocyte Dsg1 through paracrine signaling. Primary human keratinocytes were treated for 24 hours with conditioned media from melanocytes (MC) or the melanoma cell lines 1341D, 1366, 3211, YUMAC and 501mel and RNA or protein was collected. RT-PCR (A) and Western blots (B) were performed for Dsg1, Ecad and Dsg3. Significantly lower mRNA and protein expression of Dsg1, but not Ecad or Dsg3 was observed. n=3. Mean +/- SEM depicted. *p<0.05; **p<0.01; ***p<0.001. One-way ANOVA. C) Melanocytes or 1341D melanoma cells were seeded with normal primary keratinocytes and grown as 3D organotypic raft co-cultures for up to 10 days. Paraffin sections were co-stained for Dsg1 or Ecad and Mel-A. Dashed line indicates basement membrane. D) Pixel intensities were determined and the ratio of distant to proximal intensities plotted. A significant decrease in Dsg1, but not Ecad, intensity proximal to the 1341D lesions was observed. Mean +/- SEM depicted. n=3. *p<0.05, Student’s t-test.

To address whether melanoma cells reduce keratinocyte Dsg1 expression in a physiologically relevant model of epidermal differentiation, keratinocytes and melanoma cells or melanocytes were co-cultured for 6 or 10 days at an air-medium interface to stimulate keratinocyte stratification and differentiation. Of the lines tested, WM1341D cells formed discrete tumors along the basement membrane that facilitated accurate quantification of Dsg1 levels in the surrounding keratinocytes. In these 3D cultures, keratinocytes adjacent to melanoma cells, but not normal melanocytes, exhibited a significant decrease in Dsg1, but not Ecad, at the cell borders consistent with what was observed in patient melanoma lesions (Fig. 2C, D). These data show that melanoma cells can downregulate Dsg1 transcription and protein in adjacent keratinocytes, and support the idea that a secreted factor is, at least in part, responsible.

### Transcriptomic analysis reveals increased activation of keratinocyte Slug and reduced keratinocyte differentiation signaling by melanoma cells

While keratinocyte Dsg1 is vital to the formation and maintenance of a functional epidermis, the mechanisms underlying its transcriptional regulation are poorly understood. To determine what pathways were altered in keratinocytes upstream and downstream of Dsg1 loss, we performed mRNA sequencing on keratinocytes that had been treated with melanoma or melanocyte conditioned media. We utilized gene set enrichment analyses (GSEA) to analyze primary keratinocytes treated with melanoma conditioned media from WM1341D or 501mel cells compared to keratinocytes treated with conditioned media from primary melanocytes, using pathway data characterizing keratinocyte differentiation states in neonatal human skin (Fig. S2A-E) (Wang et al., 2020).

This analysis revealed a negative enrichment of gene sets characterizing the most differentiated keratinocyte states (granular and spinous) with some positive enrichment of more basal like transcriptional states (Fig. S2D, E). We previously showed that Dsg1 promotes keratinocyte differentiation through suppression of MAPK signaling. Consistent with this, upstream regulator analysis of our gene expression dataset using Ingenuity Pathway Analysis (IPA), predicted a significant increase in activation of MAPK signaling as indicated by increased ERK1/2 activation (Fig. S2F). This prediction is consistent with our previous finding that activation of MAPK signaling occurs downstream of Dsg1 loss to reduce keratinocyte differentiation.

The question remained as to what upstream factors stimulated by treatment with melanoma conditioned media might regulate Dsg1 and the associated differentiation pathways. In this regard, another predicted upstream regulator that emerged was *Snai2* (Slug), which has been associated with stem like properties of epidermal cells, as well as RAC1, which can activate Slug through PAK1 (Thaper et al., 2017; Yao et al., 2020). Slug can directly inhibit the expression of the transcription factor Grhl1, a transcriptional activator of Dsg1 and could mediate the negative enrichment of differentiated keratinocyte gene expression patterns (Mistry et al., 2014; Wilanowski et al., 2008). A potential role for keratinocyte Grhl1 in driving a permissive environment for melanoma progression has never been addressed.

### Loss of Grhl1 and increased Slug are closely tied with melanoma mediated Dsg1 loss

As Slug/Grhl1 signaling emerged as candidate regulators of Dsg1 in the context of melanoma paracrine signaling, we assessed Grhl1 mRNA and protein levels in keratinocytes treated with melanoma cell conditioned media compared to those treated with melanocyte conditioned media. Grhl1 mRNA and protein were downregulated by melanoma conditioned media but remained unchanged in Dsg1-deficient keratinocytes compared to control indicating that its loss is likely occurring upstream of melanoma induced Dsg1 downregulation (Fig. 3A-D). We also assessed the extent to which Grhl1 is decreased in keratinocytes adjacent to melanoma cells in 3D epidermal equivalent cultures (Fig. 3E). As might be predicted from previous observations that Grhl1 is associated with the epidermal differentiation program, nuclear Grhl1 was present in a gradient in control co-cultures, with the most intense nuclear staining present in the superficial, differentiated layers. Whereas Grhl1 was present in the most superficial layers of WM1341D co-cultures, expression was significantly reduced in keratinocytes surrounding tumor nests compared with control (Fig. 3E). Similarly, keratinocytes surrounding melanoma cells in patient samples exhibited reduced Grhl1 staining compared with those surrounding melanocytes in benign nevi (Fig. 3F). Grhl1 levels remained unchanged in Dsg1-deficient cultures compared to control in the 3D culture setting, further indicating that its loss likely occurs upstream of Dsg1 downregulation (Fig. 3G). We next examined melanoma regulation of keratinocyte Slug using staining in 3D cultures. Slug exhibited a basal staining pattern in melanocyte containing cultures, as is seen in normal human epidermal tissues. However, nuclear Slug levels were increased in suprabasal keratinocytes adjacent to WM1341D cells (Fig. 3H). Consistent with the decrease in Grhl1 staining in melanoma adjacent keratinocytes, Slug staining was increased in lesional keratinocytes in clinical melanoma tumor sections (Fig. 3I). We found no increase in suprabasal Slug staining in Dsg1 knockdown cultures compared to control, indicating that, like Grhl1, Slug is regulated by melanoma cells upstream of Dsg1 loss (Fig. 3J).

**Figure 3:**
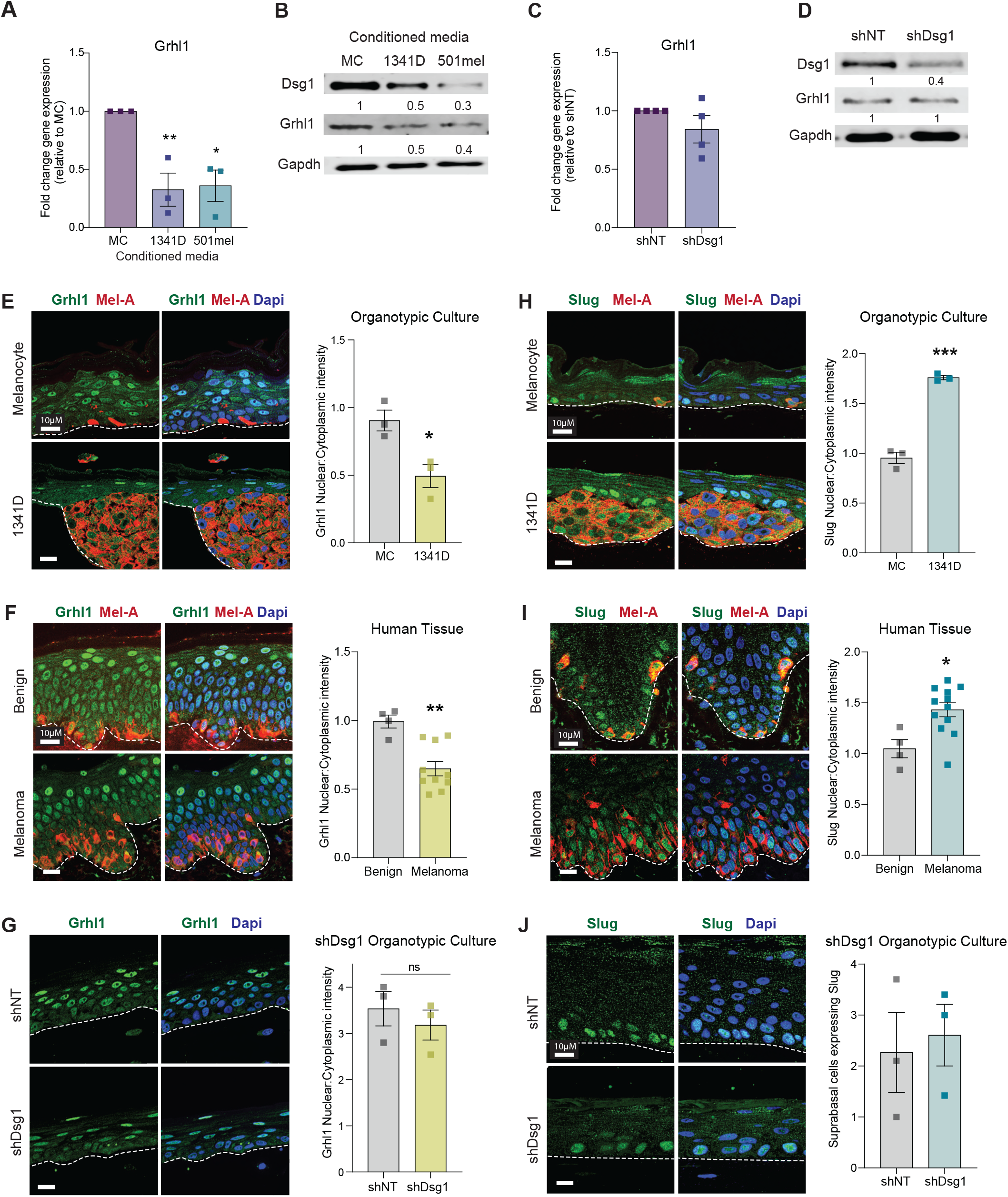
Loss of Grhl1 is associated with melanoma mediated Dsg1 loss. A-B) Primary human keratinocytes were treated for 24 hours with conditioned media from melanocytes (MC) or the melanoma cell lines 1341D and 501mel and RNA or protein was collected. RT-PCR and Western blots were performed for Grhl1. Significantly lower mRNA and protein expression of Grhl1 was observed in the keratinocytes treated with melanoma conditioned media when compared with melanocyte control conditioned media. RNA levels and quantification for blots represent average fold change. Mean +/- SEM depicted. n=3. *p<0.05, **p<0.01, *** p <0.001. one-way ANOVA. C-D) Retroviral transduction of shNT (non-targeting control) or shDsg1 knockdown vectors was performed in primary human keratinocytes and RNA or protein collected for RT-PCR and western blot validation of knockdown. No significant difference in Grhl1 mRNA expression or protein level was observed in response to Dsg1 knockdown. E) Melanocytes or 1341D melanoma cells were seeded with normal primary keratinocytes and grown as 3D organotypic raft co-cultures for 6 days. Paraffin sections were co-stained for Grhl1 and Mel-A. Pixel intensities were determined and the ratio of distant to proximal intensities plotted. A significant decrease in Grhl1 intensity proximal to the 1341D lesions was observed. Mean +/- SEM *p<0.05. Student’s t-test. F) Paraffin-embedded sections of benign nevi and melanomas were co-stained for Grhl1 and Mel-A and nuclear and cyto-plasmic pixel intensities measured in cells proximal to Mel-A stained cells and plotted as nuclear/cytoplasmic ratios. A significant decrease in keratinocyte Grhl1 intensity was observed in keratinocytes adjacent to melanoma compared with those in benign nevi. Mean +/- SEM depicted. n>3. **p<0.01. Student’s t-test. G) Similarly, 3D organotypic raft cultures comprised of shNT or shDsg1 expressing keratinocytes were grown for 6 days and stained for Grhl1. Nuclear to cytoplasmic staining intensity of Grhl1 was not statistically different between the two cultures. Mean +/- SEM depicted. n=3, Student’s t-test. H) Melanocytes or 1341D melanoma cells were seeded with primary keratinocytes and grown as 3D organotypic raft co-cultures for 6 days. Paraffin sections were co-stained for Slug and Mel-A. Pixel intensities were determined and the ratio of distant to proximal intensities plotted. A significant increase in Slug intensity proximal to the 1341D lesions was observed. +/- SEM ***p<0.001. Student’s t-test. I) Paraffin-embedded sections of benign nevi and melanomas were co-stained for Slug and Mel-A. Nuclear and cytoplasmic pixel intensities were measured in cells proximal to Mel-A stained cells and the nuclear/cytoplasmic ratios were plotted. A significant increase in keratinocyte Slug intensity was observed in keratinocytes adjacent to melanoma compared with those in benign nevi. Mean +/- SEM depicted. n>3. *p<0.05. Student’s t-test J) 3D organotypic raft cultures comprised of shNT or shDsg1 expressing keratinocytes were grown for 6 days and stained for Slug and the number of suprabasal Slug expressing cells was counted. The number of suprabasal Slug expressing cells was not different between the two cultures. n=3. Student’s t-test.

Building upon these findings, analysis of a previously published dataset (Mistry et al., 2014) revealed that loss of Slug in primary human keratinocytes led to an increase in Dsg1 and Grhl1 gene expression with no change in Ecad, similar to what we see in keratinocytes in melanoma conditioned media (Fig. S3A). Furthermore, knockout of Slug in keratinocytes using siRNA leads to a concurrent increase in Dsg1 protein levels, supporting the role of Slug as a potential mediator of Dsg1 loss in melanoma associated keratinocytes (Fig. S3B). Together these data support a model whereby melanoma cells mediate an increase in Slug and an accompanying decrease in Grhl1 and its downstream transcriptional target Dsg1 through paracrine signaling to keratinocytes.

### Loss of keratinocyte Dsg1 promotes melanoma cell migration *in vitro* and is associated with intra-epidermal movement of melanoma cells *in vivo*

We next sought to identify the biological consequences of the loss of keratinocyte Dsg1 related to malignant behavior in melanoma cells. We performed RNA sequencing of melanoma cells treated with media from control or shDsg1 expressing keratinocytes followed by GSEA using previously published melanoma differentiation state signatures (Tsoi et al., 2018). This analysis revealed an enrichment in a neural crest like signature in melanoma cells treated with Dsg1-deficient keratinocyte media (Fig. 4A). We also observed a negative enrichment of the melanocytic signature, suggesting a general pattern of de-differentiation in these cells. Consistent with this, Gene Ontology (GO) pathway analysis revealed changes in mesoderm formation, negative regulation of cell differentiation, and regulation of EMT pathways, with increased expression of genes that have been linked to neural crest formation (Fig. 4B). Given that these signatures are associated with cell movement and migration, and that loss of keratinocyte Dsg1 leads to aberrant, pagetoid movement of melanocytes in 3D epidermal equivalents, we asked whether loss of Dsg1 in keratinocytes in turn promotes melanoma cell migration.

**Figure 4:**
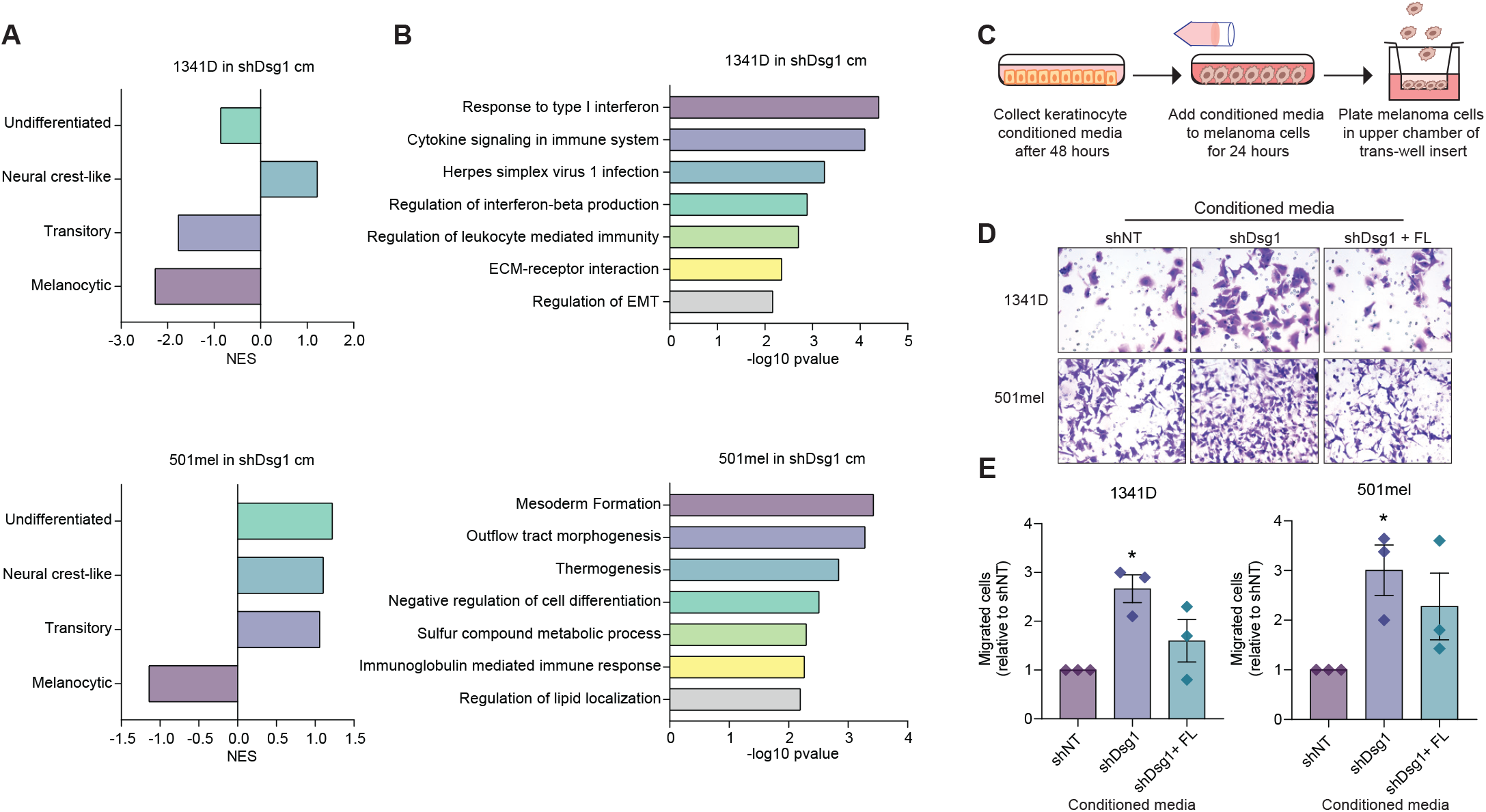
Loss of keratinocyte Dsg1 drives melanoma cell migration. Melanoma cells were cultured for 24 hours in conditioned media from keratinocytes transduced with shNT or shDsg1 knockdown vectors and collected for RNA-sequencing. A, B) Gene set enrichment analysis (GSEA) comparing differentially expressed gene sets to published signatures of melanoma signaling and differentiation. Normalized enrichment score (NES). B) Top Gene Ontology (GO) Biological Process terms overrepresented in genes upregulated by shDsg1 keratinocyte conditioned media. C) Schematic of trans-well migration assay. Melanoma cells were cultured in conditioned media from shNT expressing keratinocytes or shDsg1 expressing keratinocytes for 24 hours. 50,000 cells were then seeded in serum free media in the upper chamber of a trans-well insert (8 μM pore size). Lower wells contained DMEM supplemented with 5% fetal bovine serum (FBS). D, E) After 24 h, migrated cells were fixed and stained for visualization. Bars represent normalized migration compared with shNT control conditioned media. Mean +/- SEM depicted. n=3 *p< 0.05 Student’s t-test.

To address this question, we cultured melanoma cells in conditioned media from keratinocytes expressing Dsg1 knockdown or control and then performed a trans-well migration assay (Fig. 4C). Melanoma cells cultured in the Dsg1-deficient keratinocyte media showed increased levels of migration while re-introduction of Dsg1 into deficient keratinocytes ameliorated this effect (Fig. 4D, E). These data support the idea that Dsg1 may be important for retaining melanoma cells in the primary tumor niche.

To investigate the relationship between keratinocyte Dsg1 and melanoma cell movement *in vivo* we evaluated the extent to which melanoma spread in human patient samples was associated with Dsg1, Ecad, Grhl1 and Slug staining levels in surrounding keratinocytes. Notably, while we did see significant changes in melanoma adjacent Dsg1, Grhl1, and Slug, we observed significant variability in the degree of change of each of these proteins. When we compared the levels of keratinocyte Dsg1 staining to the number of melanoma cells which had begun to move throughout the epidermis, we observed a significant negative correlation between keratinocyte Dsg1 and melanoma epidermal spread (Fig. 5 A, B). Keratinocyte Dsg1 was not correlated with the total number of melanoma cells in the lesions, indicating that the observed change in number of migrating cells are likely not a result of increased cell number (Fig. 5C). Consistent with this, there was no correlation between number of melanoma cells in the niche and the number of cells moving throughout the epidermis.

**Figure 5.**
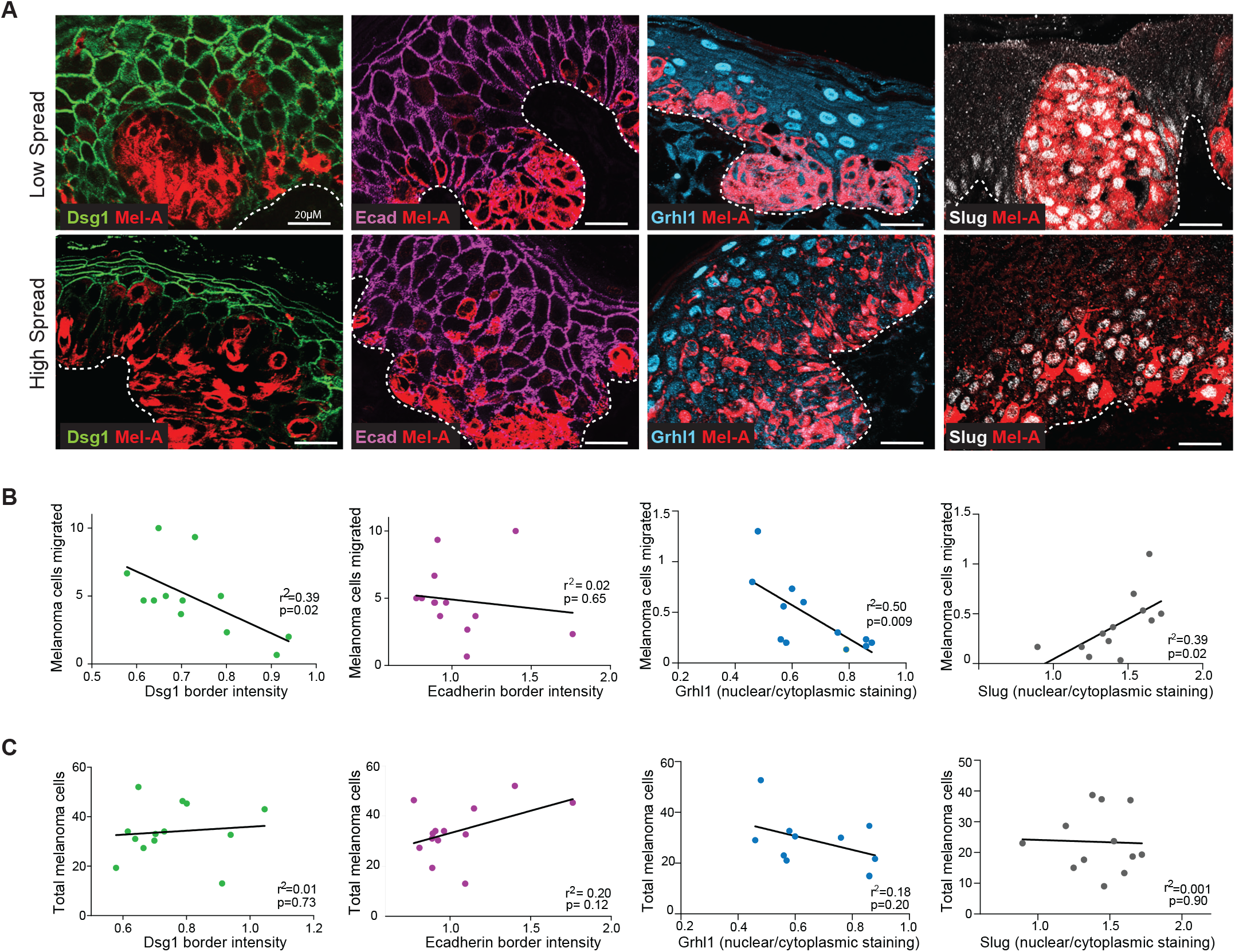
Dsg1 and its regulators are associated with epidermal spread of melanoma cells *in vivo*. Paraffin-embedded sections of melanoma lesions were stained for Dsg1, Grhl1, Slug, or Ecad and Mel-A. The number of individual melanoma cells spreading away from the basal layer, (B) as well as the total number of melanoma cells in each section (C) were counted, and then correlated to the staining intensity of Dsg1, Grhl1, Slug and Ecad. n>10. Simple linear regression.

A significant reciprocal association between Grhl1 and local melanoma spread was also observed, while Slug showed a significant positive correlation (Fig. 5 A-C). While there is also variability in keratinocyte Ecad levels in melanoma lesions, we found no significant relationship between Ecad levels and melanoma cell number or melanoma cell movement. These data are consistent with the idea that loss of keratinocyte Dsg1 and its upstream regulator Grhl1 play a role in the initial steps of melanoma cell movement and metastasis. Our data further suggest that paracrine signaling from melanoma cells may control the Dsg1/Grhl1 axis through activation of the repressor of keratinocyte differentiation gene Slug.

### Melanoma cells increase ERK1/2 dependent keratinocyte production of CXCL1 downstream of Dsg1 loss to increase melanoma cell migration

We next investigated how loss of Dsg1 in keratinocytes potentiates melanoma cell movement through paracrine signaling. Our previous work demonstrated that Dsg1-deficient keratinocytes increase secretion of inflammatory cytokines and chemokines, which are known to contribute to the formation and maintenance of the melanoma tumor niche. Chemokines previously identified as increased in the secretome of Dsg1-deficient keratinocytes include the GRO isoforms (CXCL1, CXCL2 and CXCL3), and IL-8 which bind the CXCR1 and 2 receptors, as well as the IL-6 cytokine (Arnette et al., 2019). To address whether keratinocytes grown in melanoma cell conditioned media exhibit increased chemokine expression like that seen with genetic knockdown of Dsg1, we assessed the gene expression levels of chemokines which bind the CXCR1 and CXCR2 receptors, as well as IL-6 in keratinocytes treated with media from WM1341D and 501mel cells. Of the secreted factors tested, we found that melanoma cells drive an increase in CXCL1 expression specifically in keratinocytes (Fig. 6A, Fig. S4).

**Figure 6.**
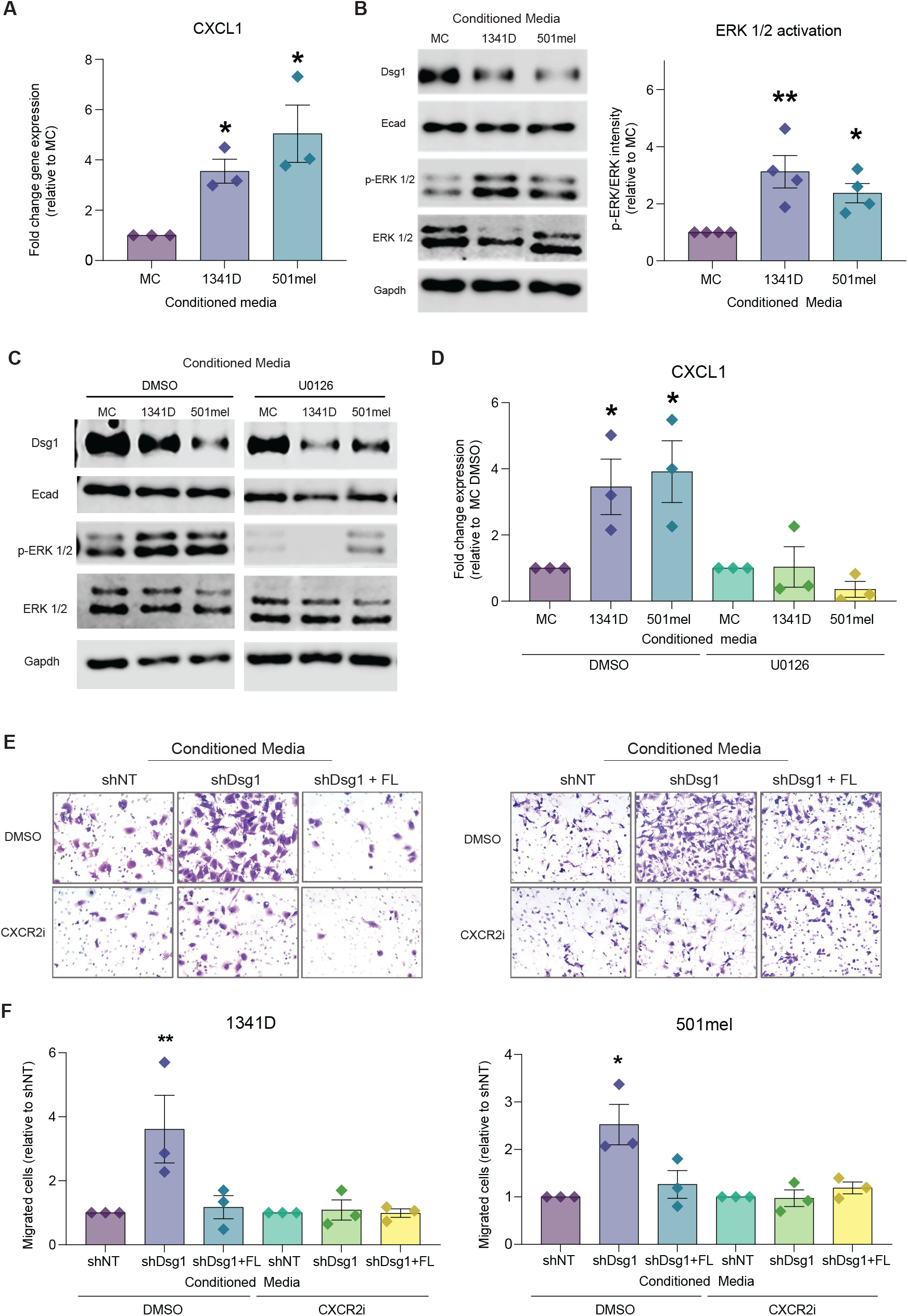
Melanoma induced loss of keratinocyte Dsg1 drives ERK1/2 dependent pro-migratory CXCL1 production. A) Primary human keratinocytes were treated for 48 hours with conditioned media from melanocytes (MC) or the melanoma cell lines 1341D and 501mel and RNA or protein was collected. RT-PCR was performed for CXCL1 expression. Mean +/- SEM depicted. n=3. *p< 0.05, One-way ANOVA. B) Western blot was performed for Dsg1, Ecad, ERK1/2 and phospho-ERK1/2 Mean +/- SEM depicted. n=4, *p< 0.05 **p<0.01. One-way ANOVA. Increased ERK1/2 phosphorylation was observed in keratinocytes treated with melanoma conditioned media. C) Primary human keratinocytes were treated with melanocyte or melanoma conditioned media with either DMSO or 5 μM U0126 (MEK1/2 inhibitor). Western blot was performed to confirm a decrease in ERK1/2 phosphorylation in U0126 treated cells. n=3. D) CXCL1 gene expression was no longer increased in keratinocytes treated with melanoma conditioned media in the presence of U0126 n=3. *p< 0.05. One-way ANOVA. E, F) Melanoma cells were treated with conditioned media from shNT, shDsg1, or shDsg1 plus a full length (FL) wild-type Dsg1 rescue expressing keratinocytes for 24 hours in the presence of DMSO or the CXCR2 inhibitor (500 nM SB22502) then plated for trans-well migration and collected after 24 hours. Loss of keratinocyte Dsg1 no longer increased melanoma cell migration when the CXCL1 receptor, CXCR2, was inhibited Mean +/- SEM depicted. n=3 *p < 0.05 **p< 0.01. One-way ANOVA.

As mentioned above, Upstream Regulator analysis predicted an increase in ERK1/2 signaling, which can increase levels of CXCL1 (Cheng et al., 2019; Schweppe et al., 2006). Our previous work showed that genetic interference with Dsg1 results in increased ERK1/2 signaling. Indeed, when active p-ERK/total ERK was measured in keratinocytes grown in melanoma cell conditioned media, a significant increase was observed, like that which occurs in cells with genetic interference of Dsg1 (Fig. 6B). To address whether the observed increase in CXCL1 depended on the ERK1/2 pathway, we treated cells in melanoma conditioned media with the MEK1/2 inhibitor U0126. U0126 effectively reduced both ERK1/2 activation as well as CXCL1 gene expression (Fig. 6C, D). Dsg1 levels are not rescued by ERK1/2 inhibition, indicating that melanoma driven ERK1/2 signaling is likely downstream of and mediated by Dsg1 loss as indicated by our transcriptomic analysis.

CXCL1 exclusively binds to the CXCR2 receptor in melanoma cells. To confirm the role of CXCL1 in melanoma cell migration we treated melanoma cells with a combination of keratinocyte conditioned media and DMSO or the CXCR2 inhibitor SB225002. When the CXCL1 receptor CXCR2 was inhibited in melanoma cells, Dsg1-deficient keratinocytes were no longer able to induce migration, demonstrating a role for specific chemokine signaling in keratinocyte modulation of melanoma cell behavior (Fig. 6E, F). Together these data highlight a novel mechanism of initiation or increase of melanoma cell migration through direct manipulation of tumor niche keratinocytes.

## Discussion

Melanoma is one of the most aggressive skin cancers, due in part to its propensity to metastasize. The composition of the tumor microenvironment has an undeniably important influence on tumor cell behavior, including metastatic progression. Much of the work done investigating melanoma metastasis has been focused on cells once they have already escaped from the epidermis (Vandyck et al., 2021). Here we examine the underappreciated role of keratinocytes in the initiation (or inhibition) of melanoma cell movement within the primary epidermal niche. Our work establishes a role for the keratinocyte-specific cadherin Dsg1 as a regulator of chemokine signaling whose loss is associated with cell spread in the melanoma tumor niche (Fig. 7). Furthermore, the studies shed light on the understudied mechanisms by which melanoma cells manipulate their epidermal microenvironment to facilitate metastatic progression.

**Figure 7.**
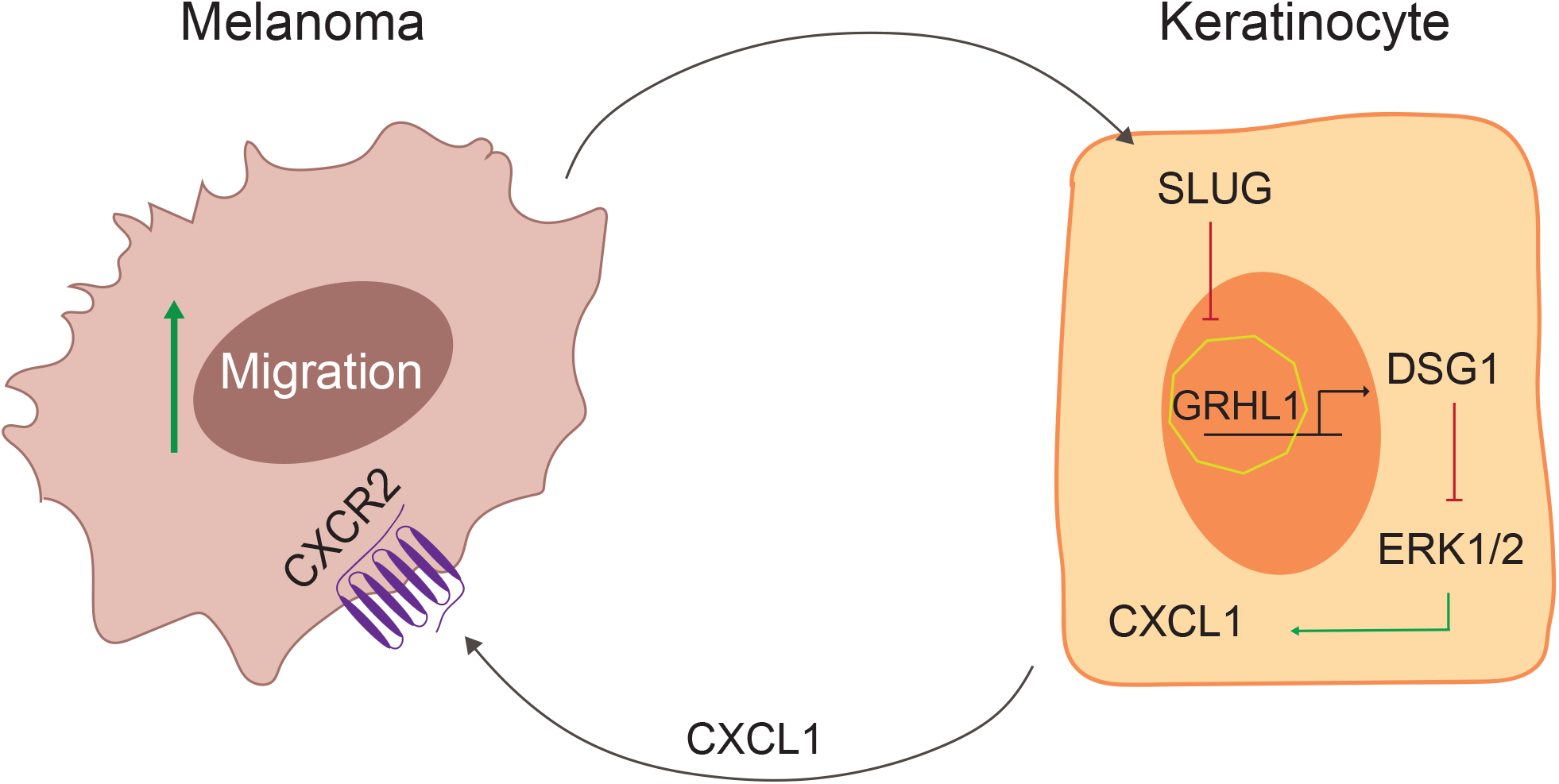
Model: Bi-directional paracrine signaling between melanoma cells and keratinocytes potentiates melanoma cell movement. Our data support a model whereby factors secreted from melanoma cells activate keratinocyte Slug, which in turn decreases Grhl1, a transcriptional activator of Dsg1. The consequent loss of Dsg1 activates ERK1/2 to increase keratinocyte CXCL1 production, which in turn promotes melanoma cell migration through activation of CXCR2.

Previous work has highlighted the importance of keratinocytes in restricting newly transforming melanocytes to their normal location in the basal epidermis. In particular, the role of cadherins in restricting the movement of melanoma cells has focused on their canonical roles in cell-cell adhesion. For instance, loss of Ecad, which is expressed in both keratinocytes and melanocytes, occurs in progressive fashion in primary/invasive melanomas (Haass et al., 2005). Re-introduction of Ecad into Ecad-deficient melanoma cells inhibited invasion into the dermis of 3D reconstructed epidermis through downregulation of invasion receptors MUC18 and β3 integrin (Hsu et al., 2000) and also impairs melanoma cell invasion in response to CXCL12 (Molina-Ortiz et al., 2009). The loss of Ecad in melanoma may also occur in concert with upregulation of pro-migratory N-cadherin, possibly resulting in a cadherin switch similar to that reported for other tumors (Siret et al., 2015). Like Ecad, P-cadherin in the tumor environment promotes keratinocyte-melanoma cell contact. However, in this case the interaction supports the expansion of melanoma cells and progression towards a more aggressive phenotype and poor survival in humans (Mescher et al., 2017). Overall, previous work on cadherins in melanoma progression has focused on the idea of permissive migration due to removal of physical barriers or tumor progression due to cell contact mediated signaling.

Here, we provide evidence for a novel paracrine function for the desmosomal cadherin, Dsg1, in shaping the early melanoma microenvironment. While playing an essential role in the physical barrier, Dsg1 has also emerged as a cell autonomous regulator of signaling pathways that complement but do not depend on its physical role in adhesion (Arnette et al., 2019; Getsios et al., 2009; Harmon et al., 2013). Dsg1 loss due to genetic or environmental stimuli revealed a role in controlling differentiation and inflammation (Godsel et al., 2022). The data presented here add to Dsg1 functions, and are consistent with a model whereby its loss in the context of melanoma drives melanoma cell migration. Changes in Dsg1 expression occur before any detectable decrease in Ecad, supporting the idea that these changes occur early in tumor development and that melanoma spread is not an indirect result of loss of this classic cadherin. Consistent with this, melanoma conditioned media is sufficient to mediate a reduction in Dsg1 mRNA and protein, in the absence of any significant changes in Ecad expression.

Importantly, we have defined for the first time a potential transcriptional regulatory pathway co-opted by melanoma cells to reduce Dsg1 expression through Grhl1 and its upstream transcriptional repressor Slug. While Grhl1 has been shown to regulate epidermal homeostasis by controlling the differentiation program and barrier function (Wilanowski et al., 2008), a role in melanoma development has not been reported. Likewise, Slug has been linked to melanoma cell metastasis but not in the creation of a pro-melanoma keratinocyte compartment (Miao et al., 2021; Shirley et al., 2012). Importantly, while increased Slug likely plays additional roles in shaping the early melanoma tumor environment, genetic targeting of Dsg1 is sufficient to initiate the signaling cascade that stimulates increased melanoma cell movement *in vitro*.

Our observation that melanoma cells treated with conditioned media from Dsg1-deficient keratinocytes exhibit a neural crest like signature is consistent with previous studies suggesting malignant melanoma cells hijack the neural crest program to promote plasticity and facilitate invasion and metastasis (Diener and Sommer, 2021; Wessely et al., 2021). We further identify a CXCL1/CXCR2 axis as important for the increase in melanoma cell movement. CXCL1 signaling has been linked to stem cell phenotypes and tumor progression and invasion in other cancers (Tang et al., 2012; Wessely et al., 2021) and CXCR2 introduction into melanoma cells increases cell proliferation, motility and invasion (Singh et al., 2009). CXCR2 may be more prevalent in late-stage melanomas and thus it will be important to explore other potential pathways in the early melanoma niche that could link Dsg1 reduction in keratinocytes to melanoma spread at the earliest stages (Sharma et al., 2010).

Finally, here we uncovered novel paracrine regulation of melanoma behavior by the keratinocyte-specific molecule Dsg1. However, this cadherin also controls cell-cell adhesion and the mechanical properties of keratinocytes that along with paracrine signaling could impact melanoma behavior. In the future, understanding how both cell contact mediated and paracrine signaling changes are integrated will be important for acquiring a cohesive understanding of factors shaping the tumor niche. Futhermore, study of how loss of Dsg1 contributes to the plasticity of the tumor cells, whether it stimulates migration of an existing rare population of melanoma cells or lowers the threshold for the emergence of a new population with neural crest features will be important for understanding the molecular mechanisms of communication between these two populations (Emert et al., 2021; Shaffer et al., 2017). As melanoma cells do not acquire new mutations (Shain et al., 2018) once they have left the epidermis, keratinocyte cross talk could be a key shaper of melanoma behavior in the epidermal tumor niche. Changes in melanoma cells caused by a Dsg1-deficient microenvironment maintained over time could be targetable.

## Materials and Methods

### Cell Culture

Keratinocytes and melanocytes were isolated from discarded neonatal foreskin provided by the Northwestern University Skin Biology and Diseases Research-Based Center (NUSBDRC) as previously described (Roth-Carter et al., 2022). Keratinocytes were grown in M154 medium supplemented with human keratinocyte growth supplement (Life Technologies), 1,000 x gentamycin/amphotericin B solution (Life Technologies), and 0.07mM CaCl_2_. Confluent keratinocyte monolayers were differentiated in the media described with addition of 1.2 mM CaCl_2_. Keratinocytes were transduced with retroviral supernatants produced from Phoenix cells (provided by G. Nolan, Stanford University, Stanford, CA) as previously described (Getsios et al., 2004; Simpson et al., 2010). Melanocytes were cultured in Opti-MEM (Life Technologies) containing 1% penicillin/streptomycin (Corning), 5% fetal bovine serum (FBS) (Sigma-Aldrich), 10 ng/ml bFGF (ConnStem Inc), 1 ng/ml heparin (Sigma-Aldrich), 0.1 mM N6, 2’-O-dibutyryladenosine 3:5-cyclic monophosphate (dbcAMP; Sigma-Aldrich), and 0.1 mM 3-isobutyl-1-methyl xanthine (IBMX; Sigma-Aldrich). Melanoma cells were cultured in DMEM containing sodium pyruvate (Life Technologies) with 1% penicillin/streptomycin, 5% FBS (Sigma-Aldrich). WM1341D, WM1366, WM3211 lines were purchased from Rockland Immunochemicals Inc. YUMAC and 501mel lines were kindly gifted by Jaehyuk Choi at Northwestern University. All cell lines were authenticated with DNA (STR) profiling and tested to rule out mycoplasma contamination.

### Organotypic skin cultures

Organotypic cultures were grown as previously described (Arnette et al., 2016; Getsios et al., 2009) with the addition of melanocytes or melanoma cells co-seeded with keratinocytes at a 1:5 ratio to mimic the ratio seen in the basal layer of normal skin (Roth-Carter et al., 2022). Organotypic cultures fixed in 10% neutral buffered formalin were embedded in paraffin blocks and cut into 4 μm sections.

### Preparation of conditioned media

Conditioned media from keratinocytes was prepared as previously described (Arnette et al., 2019). Melanocytes or melanoma cells were grown to 70-80% confluence, at which point the media was refreshed and then collected after 72 hours. Conditioned media was added to keratinocytes immediately upon collection.

### Antibodies and reagents

The mouse monoclonal antibodies used were: P124 (anti-Dsg1 extracellular domain; Progen, Heidelberg, Germany), anti-Dsg3 (Progen) 27B2 (anti-Dsg1 cytodomain; Life Technologies), and HMB45 (anti-Melanoma gp100; Thermo Fisher Scientific), total ERK1/2 (Cell Signaling). The rabbit monoclonal antibody EP1576Y (ab52642, anti-S100 beta; Abcam,) was used. Rabbit polyclonal antibodies used were: HECD1 (anti-E-cadherin; Takara), anti-Melan-A (ab15468; Abcam), anti-GAPDH (G9545, glyceraldehyde-3-phosphate dehydrogenase; Sigma-Aldrich), anti-GRHL1 (Thermo Fisher), anti-Slug (Cell Signaling), anti-p-ERK (Cell Signaling), and anti-ERK (Cell Signaling). Secondary antibodies for immunoblotting were goat anti-mouse and goat anti-rabbit (LI-COR). Secondary antibodies for immunofluorescence were goat anti-mouse and goat-anti-rabbit linked to fluorophores of 488, 568, and 647 nm (Alexa Fluor; Life Technologies). DAPI was used to stain nuclei. The MEK1/2 inhibitor (U0126) was purchased from Cell Signaling and used at a 5 μM final concentration. The CXCR2 inhibitor (SB22502) was purchased from Selleck Chemicals and used at a 500 nM final concentration.

### DNA constructs

LZRS-shDsg1 (Dsg1shRNA) and LZRS-Flag Dsg1 were generated as described (Getsios et al., 2009; Simpson et al., 2010). LZRS – NTshRNA was generated as described (Arnette et al., 2016; Getsios et al., 2009). Human SNAI2 siRNA SMART pool was purchased from Dharmacon (5’- UCUCUCCUCUUUCCGGAUA-3’, 5’-GCGAUGCCCAGUCUAGAAA-3’, 5’- ACAGCGAACUGGACACACA-3’, 5’-GAAUGUCUCUCCUGCACAA-3’) and transfected as previously described (Arnette et al., 2019).

### Trans-well migration assay

50,000 cells were seeded in 500 μL phenol red-free Opti-MEM in the upper chamber of a 24-well trans-well chamber. DMEM containing 5% FBS was used as a chemoattractant in the lower wells. After 24 hours, inner membranes were scrubbed to remove non-migrated cells. Cells on the outer membranes were fixed in formalin and stained with 0.1% crystal violet. Membranes were excised from the trans-well insert and mounted on glass slides. Migrated cells were visualized by microscopy and counted.

### qPCR

Total RNA was extracted from cells using the Quick-RNA miniprep kit (Zymo Research) following the manufacturer’s protocol. CDNA was synthesized using 1 μg of RNA using the Superscript III First Strand Synthesis Kit (Thermo Fisher). Quantitative PCR was performed on the QuantStudio 3 instrument (Thermo Fisher), using SYBR Green PCR master mix (Thermo Fisher). Relative mRNA levels were calculated using the ΔΔCT method normalized to GAPDH.

### Immunofluorescence

For immunostaining, tissue sections were baked at 60°C overnight and de-paraffinized using xylenes. Samples were then rehydrated through a series of ethanol and PBS washes. Tissues were permeabilized in 0.5% Triton X-100 in PBS. Antigen retrieval was performed by incubation in 0.01 M citrate buffer at 95°C for 15 minutes. Sections were blocked in blocking buffer (1% BSA, 2% normal goat serum in PBS) for 60 minutes at 37°C. Samples were then incubated in primary antibody at 4°C overnight, followed by incubation in secondary antibody for 1 hour at 37°C. Images were acquired using an AxioVison Z1 system (Carl Zeiss) with Apotome slide module, an AxioCam MRm digital camera, and either a 20x (0.8 NA Plan-Apochromat) or 40x (1.4 NA, Plan-Apochromat) oil objective. Image analysis was carried out using ImageJ software.

### Immunoblot

Whole cell lysates were collected from confluent monolayers in urea-SDS buffer (8 M urea/1% SDS/60 mM Tris (pH 6.8)/5% ß-mercaptoethanol/10% glycerol) and sonicated. Samples separated by SDS-PAGE were transferred to nitrocellulose, blocked in Odyssey Blocking Buffer (LI-COR Biosciences), and incubated with primary antibodies in blocking buffer overnight at 4°C. After a series of PBS washes, secondary antibodies diluted in blocking buffer at 1:10,000 concentration were added to blots for 1 hour at room temperature then washed with a series of PBS washes. Protein bands were imaged using LI-COR Odyssey FC (LI-COR Biosciences). Densitometric analysis was performed on scanned blots using LI-COR Image studio software.

### RNA sequencing

Differentiated keratinocytes were cultured with conditioned media for 48 hours and melanoma cells were cultured with conditioned media from keratinocytes expressing shNT or shDsg1 for 24 hours. Total RNA was extracted from cells using the Quick-RNA miniprep kit (Zymo Research) following the manufacturer’s protocol. Novogene Co., LTD conducted preparation of the mRNA library and transcriptome sequencing. Analyses were performed in R (version 4.1.2) using Bioconductor libraries and R statistical packages. Differential expression analysis was performed using DESeq2. Genes with p-value < 0.05 and |log2(FoldChange)| > 1 were considered to be differentially expressed. Pathway analysis was performed using Ingenuity Pathway Analysis (IPA) for Upstream Regulators and GSEA.

### Microarray analysis

Microarray analyses of SNAI2 knockdown keratinocytes (GSE55269) were performed in R (version 4.1.2) using Bioconductor libraries and R statistical packages. Affymetrix probe IDs were converted to unique Entrez IDs for analysis, and differential expression comparisons were performed using the Limma package.

### Statistical analysis

Statistical analysis was performed using GraphPad Prism 8.0 software. All experiments were performed at least three times. Measured data were represented as the mean +/- SEM. One-way analysis of variance (ANOVA) or two-tail Student’s t-test were applied to compare quantitative data. P-values for each analysis are marked on figures, and the level of statistical significance was defined as *p<0.05, ** p<0.01, ***p<0.001, ****p<0.0001.

## ACKNOWLEDGMENTS

Research was supported by NIH/NIAMS P30 AR075049 awarded to Northwestern University Skin Biology & Diseases Resource-Based Center. Histology services were provided by the Northwestern University Mouse Histology and Phenotyping Laboratory, which is supported by NCI P30-CA060553, awarded to the Robert H Lurie Comprehensive Cancer Center. This work was supported by NIH/NCI R01 CA228196 with additional support from NIH/NIAMS R01 AR041836, NIH/NIAMS R01 AR043380, a Leo Foundation Grant and the J.L. Mayberry Endowment to KJG. JLP was supported by NIH/NCI T32 CA009560. HEB was supported by NIH/NCI T32 CA080621-14. QRRC was supported by NIH/NCI T32 CA070085. JTS was supported by NIH/NIAMS P30-AR069589.

## AUTHOR CONTRIBUTIONS

KJG, HEB and CRA conceived of and designed experiments. HEB, JLK and JLP acquired data. HEB, QRRC and JLP performed imaging and data analysis. CRA and JLJ provided training and technical assistance. PG and JTS provided tissue samples. KJG and HEB wrote the manuscript.

## DATA AVAILABILITY

The dataset used during the study is available from the corresponding author upon request.

## REFERENCES

Arnette, C., J.L. Koetsier, P. Hoover, S. Getsios, and K.J. Green. 2016. In Vitro Model of the Epidermis: Connecting Protein Function to 3D Structure. Methods Enzymol. 569:287–308.

Arnette, C.R., Q.R. Roth-Carter, J.L. Koetsier, J.A. Broussard, H.E. Burks, K. Cheng, C. Amadi, P. Gerami, J.L. Johnson, and K.J. Green. 2019. Keratinocyte cadherin desmoglein 1 controls melanocyte behavior through paracrine signaling. Pigment Cell Melanoma Res.

Brandner, J.M., and N.K. Haass. 2013. Melanoma’s connections to the tumour microenvironment. Pathology. 45:443–452.

Cheng, Y., X.L. Ma, Y.Q. Wei, and X.W. Wei. 2019. Potential roles and targeted therapy of the CXCLs/CXCR2 axis in cancer and inflammatory diseases. Biochim Biophys Acta Rev Cancer. 1871:289–312.

Diener, J., and L. Sommer. 2021. Reemergence of neural crest stem cell-like states in melanoma during disease progression and treatment. Stem Cells Transl Med. 10:522–533.

Emert, B.L., C.J. Cote, E.A. Torre, I.P. Dardani, C.L. Jiang, N. Jain, S.M. Shaffer, and A. Raj. 2021. Variability within rare cell states enables multiple paths toward drug resistance. Nat Biotechnol. 39:865–876.

Galbo, P.M., Jr., X. Zang, and D. Zheng. 2021. Molecular Features of Cancer-associated Fibroblast Subtypes and their Implication on Cancer Pathogenesis, Prognosis, and Immunotherapy Resistance. Clin Cancer Res. 27:2636–2647.

Getsios, S., E.V. Amargo, R.L. Dusek, K. Ishii, L. Sheu, L.M. Godsel, and K.J. Green. 2004. Coordinated expression of desmoglein 1 and desmocollin 1 regulates intercellular adhesion. Differentiation. 72:419–433.

Getsios, S., C.L. Simpson, S. Kojima, R. Harmon, L.J. Sheu, R.L. Dusek, M. Cornwell, and K.J. Green. 2009. Desmoglein 1-dependent suppression of EGFR signaling promotes epidermal differentiation and morphogenesis. J Cell Biol. 185:1243–1258.

Godsel, L.M., Q.R. Roth-Carter, J.L. Koetsier, L.C. Tsoi, A.L. Huffine, J.A. Broussard, G.N. Fitz, S.M. Lloyd, J. Kweon, H.E. Burks, M. Hegazy, S. Amagai, P.W. Harms, X. Xing, J. Kirma, J.L. Johnson, G. Urciuoli, L.T. Doglio, W.R. Swindell, R. Awatramani, E. Sprecher, X. Bao, E. Cohen-Barak, C. Missero, J.E. Gudjonsson, and K.J. Green. 2022. Translational implications of Th17-skewed inflammation due to genetic deficiency of a cadherin stress sensor. J Clin Invest. 132.

Golan, T., A.R. Messer, A. Amitai-Lange, Z. Melamed, R. Ohana, R.E. Bell, O. Kapitansky, G. Lerman, S. Greenberger, M. Khaled, N. Amar, J. Albrengues, C. Gaggioli, P. Gonen, Y. Tabach, D. Sprinzak, R. Shalom-Feuerstein, and C. Levy. 2015. Interactions of Melanoma Cells with Distal Keratinocytes Trigger Metastasis via Notch Signaling Inhibition of MITF. Mol Cell. 59:664–676.

Green, K.J., Q.R. Roth-Carter, C.M. Niessen, and S.A. Nichols. 2020. Tracing the Origins of the Desmosome: A Vertebrate Innovation. Curr Biol. 30:R535–R543.

Haass, N.K., K.S. Smalley, L. Li, and M. Herlyn. 2005. Adhesion, migration and communication in melanocytes and melanoma. Pigment cell research / sponsored by the European Society for Pigment Cell Research and the International Pigment Cell Society. 18:150–159.

Harmon, R.M., C.L. Simpson, J.L. Johnson, J.L. Koetsier, A.D. Dubash, N.A. Najor, O. Sarig, E. Sprecher, and K.J. Green. 2013. Desmoglein-1/Erbin interaction suppresses ERK activation to support epidermal differentiation. J Clin Invest. 123:1556–1570.

Hegazy, M., A.L. Perl, S.A. Svoboda, and K.J. Green. 2022. Desmosomal Cadherins in Health and Disease. Annu Rev Pathol. 17:47–72.

Hsu, M.Y., F.E. Meier, M. Nesbit, J.Y. Hsu, P. Van Belle, D.E. Elder, and M. Herlyn. 2000. E-cadherin expression in melanoma cells restores keratinocyte-mediated growth control and down-regulates expression of invasion-related adhesion receptors. Am J Pathol. 156:1515–1525.

Johnson, J.L., J.L. Koetsier, A. Sirico, A.T. Agidi, D. Antonini, C. Missero, and K.J. Green. 2014. The desmosomal protein desmoglein 1 aids recovery of epidermal differentiation after acute UV light exposure. J Invest Dermatol. 134:2154–2162.

Kalaora, S., A. Nagler, J.A. Wargo, and Y. Samuels. 2022. Mechanisms of immune activation and regulation: lessons from melanoma. Nat Rev Cancer. 22:195–207.

Kowalczyk, A.P., and K.J. Green. 2013. Structure, function, and regulation of desmosomes. Prog Mol Biol Transl Sci. 116:95–118.

Mescher, M., P. Jeong, S.K. Knapp, M. Rubsam, M. Saynisch, M. Kranen, J. Landsberg, M. Schlaak, C. Mauch, T. Tuting, C.M. Niessen, and S. Iden. 2017. The epidermal polarity protein Par3 is a non-cell autonomous suppressor of malignant melanoma. J Exp Med.

Miao, Y., W. Zhang, S. Liu, X. Leng, C. Hu, and H. Sun. 2021. HOXC10 promotes growth and migration of melanoma by regulating Slug to activate the YAP/TAZ signaling pathway. Discov Oncol. 12:12.

Mistry, D.S., Y. Chen, Y. Wang, K. Zhang, and G.L. Sen. 2014. SNAI2 controls the undifferentiated state of human epidermal progenitor cells. Stem Cells. 32:3209–3218.

Molina-Ortiz, I., R.A. Bartolome, P. Hernandez-Varas, G.P. Colo, and J. Teixido. 2009. Overexpression of E-cadherin on melanoma cells inhibits chemokine-promoted invasion involving p190RhoGAP/p120ctn-dependent inactivation of RhoA. J Biol Chem. 284:15147–15157.

Polivka, L., S. Hadj-Rabia, E. Bal, S. Leclerc-Mercier, M. Madrange, Y. Hamel, D. Bonnet, S. Mallet, H. Lepidi, C. Ovaert, P. Barbet, C. Dupont, B. Neven, A. Munnich, L.M. Godsel, F. Campeotto, R. Weil, E. Laplantine, S. Marchetto, J.P. Borg, W.I. Weis, J.L. Casanova, A. Puel, K.J. Green, C. Bodemer, and A. Smahi. 2018. Epithelial barrier dysfunction in desmoglein-1 deficiency. J Allergy Clin Immunol. 142:702–706 e707.

Roth-Carter, Q.R., J.L. Koetsier, J.A. Broussard, and K.J. Green. 2022. Organotypic Human Skin Cultures Incorporating Primary Melanocytes. Curr Protoc. 2:e536.

Samuelov, L., O. Sarig, R.M. Harmon, D. Rapaport, A. Ishida-Yamamoto, O. Isakov, J.L. Koetsier, A. Gat, I. Goldberg, R. Bergman, R. Spiegel, O. Eytan, S. Geller, S. Peleg, N. Shomron, C.S. Goh, N.J. Wilson, F.J. Smith, E. Pohler, M.A. Simpson, W.H. McLean, A.D. Irvine, M. Horowitz, J.A. McGrath, K.J. Green, and E. Sprecher. 2013. Desmoglein 1 deficiency results in severe dermatitis, multiple allergies and metabolic wasting. Nat Genet. 45:1244–1248.

Schweppe, R.E., T.H. Cheung, and N.G. Ahn. 2006. Global gene expression analysis of ERK5 and ERK1/2 signaling reveals a role for HIF-1 in ERK5-mediated responses. J Biol Chem. 281:20993–21003.

Shaffer, S.M., M.C. Dunagin, S.R. Torborg, E.A. Torre, B. Emert, C. Krepler, M. Beqiri, K. Sproesser, P.A. Brafford, M. Xiao, E. Eggan, I.N. Anastopoulos, C.A. Vargas-Garcia, A. Singh, K.L. Nathanson, M. Herlyn, and A. Raj. 2017. Rare cell variability and drug-induced reprogramming as a mode of cancer drug resistance. Nature. 546:431–435.

Shain, A.H., N.M. Joseph, R. Yu, J. Benhamida, S. Liu, T. Prow, B. Ruben, J. North, L. Pincus, I. Yeh, R. Judson, and B.C. Bastian. 2018. Genomic and Transcriptomic Analysis Reveals Incremental Disruption of Key Signaling Pathways during Melanoma Evolution. Cancer Cell. 34:45–55 e44.

Sharma, B., S. Singh, M.L. Varney, and R.K. Singh. 2010. Targeting CXCR1/CXCR2 receptor antagonism in malignant melanoma. Expert Opin Ther Targets. 14:435–442.

Shirley, S.H., V.R. Greene, L.M. Duncan, C.A. Torres Cabala, E.A. Grimm, and D.F. Kusewitt. 2012. Slug expression during melanoma progression. Am J Pathol. 180:2479–2489.

Simpson, C.L., S. Kojima, and S. Getsios. 2010. RNA interference in keratinocytes and an organotypic model of human epidermis. Methods Mol Biol. 585:127–146.

Singh, S., K.C. Nannuru, A. Sadanandam, M.L. Varney, and R.K. Singh. 2009. CXCR1 and CXCR2 enhances human melanoma tumourigenesis, growth and invasion. Br J Cancer. 100:1638–1646.

Siret, C., C. Terciolo, A. Dobric, M.C. Habib, S. Germain, R. Bonnier, D. Lombardo, V. Rigot, and F. Andre. 2015. Interplay between cadherins and alpha2beta1 integrin differentially regulates melanoma cell invasion. Br J Cancer. 113:1445–1453.

Tang, K.H., S. Ma, T.K. Lee, Y.P. Chan, P.S. Kwan, C.M. Tong, I.O. Ng, K. Man, K.F. To, P.B. Lai, C.M. Lo, X.Y. Guan, and K.W. Chan. 2012. CD133(+) liver tumor-initiating cells promote tumor angiogenesis, growth, and self-renewal through neurotensin/interleukin-8/CXCL1 signaling. Hepatology. 55:807–820.

Thaper, D., S. Vahid, K.M. Nip, I. Moskalev, X. Shan, S. Frees, M.E. Roberts, K. Ketola, K.W. Harder, C. Gregory-Evans, J.L. Bishop, and A. Zoubeidi. 2017. Targeting Lyn regulates Snail family shuttling and inhibits metastasis. Oncogene. 36:3964–3975.

Tsoi, J., L. Robert, K. Paraiso, C. Galvan, K.M. Sheu, J. Lay, D.J.L. Wong, M. Atefi, R. Shirazi, X. Wang, D. Braas, C.S. Grasso, N. Palaskas, A. Ribas, and T.G. Graeber. 2018. Multi-stage Differentiation Defines Melanoma Subtypes with Differential Vulnerability to Drug-Induced Iron-Dependent Oxidative Stress. Cancer Cell. 33:890–904 e895.

Upadhyay, P.R., T. Ho, and Z.A. Abdel-Malek. 2021. Participation of keratinocyte- and fibroblast-derived factors in melanocyte homeostasis, the response to UV, and pigmentary disorders. Pigment Cell Melanoma Res. 34:762–776.

Vandyck, H.H., L.M. Hillen, F.M. Bosisio, J. van den Oord, A. Zur Hausen, and V. Winnepenninckx. 2021. Rethinking the biology of metastatic melanoma: a holistic approach. Cancer Metastasis Rev. 40:603–624.

Villanueva, J., and M. Herlyn. 2008. Melanoma and the tumor microenvironment. Curr Oncol Rep. 10:439–446.

Wang, J.X., M. Fukunaga-Kalabis, and M. Herlyn. 2016. Crosstalk in skin: melanocytes, keratinocytes, stem cells, and melanoma. J Cell Commun Signal. 10:191–196.

Wang, S., M.L. Drummond, C.F. Guerrero-Juarez, E. Tarapore, A.L. MacLean, A.R. Stabell, S.C. Wu, G. Gutierrez, B.T. That, C.A. Benavente, Q. Nie, and S.X. Atwood. 2020. Single cell transcriptomics of human epidermis identifies basal stem cell transition states. Nat Commun. 11:4239.

Wessely, A., T. Steeb, C. Berking, and M.V. Heppt. 2021. How Neural Crest Transcription Factors Contribute to Melanoma Heterogeneity, Cellular Plasticity, and Treatment Resistance. Int J Mol Sci. 22.

Wilanowski, T., J. Caddy, S.B. Ting, N.R. Hislop, L. Cerruti, A. Auden, L.L. Zhao, S. Asquith, S. Ellis, R. Sinclair, J.M. Cunningham, and S.M. Jane. 2008. Perturbed desmosomal cadherin expression in grainy head-like 1-null mice. Embo J. 27:886–897.

Yao, D., C. Li, M.S.R. Rajoka, Z. He, J. Huang, J. Wang, and J. Zhang. 2020. P21-Activated Kinase 1: Emerging biological functions and potential therapeutic targets in Cancer. Theranostics. 10:9741–9766.

Yardman-Frank, J.M., and D.E. Fisher. 2021. Skin pigmentation and its control: From ultraviolet radiation to stem cells. Exp Dermatol. 30:560–571.

