## Supplemental figures for "Keratinocyte desmosomal cadherin Desmoglein 1 as a mediator and target of paracrine signaling in the melanoma niche"

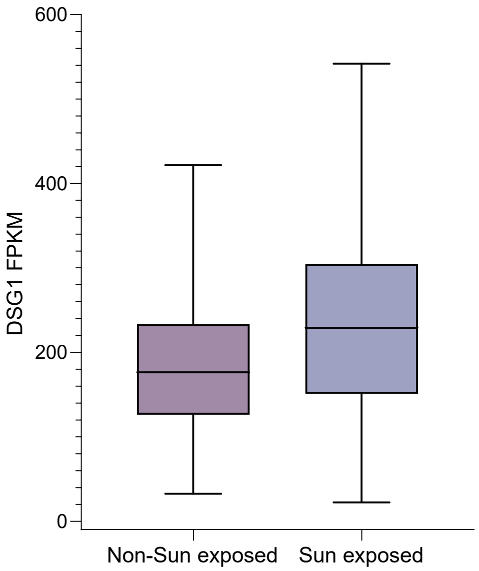


**Supplementary Figure 1. GTEx (Genotype-Tissue Expression) data showing gene expression of Dsg1 in sun-exposed and non-sun exposed skin.** Dsg1 mRNA levels were not altered in chronically sun-exposed skin. FPKM, bars represent minimum and maximum values.

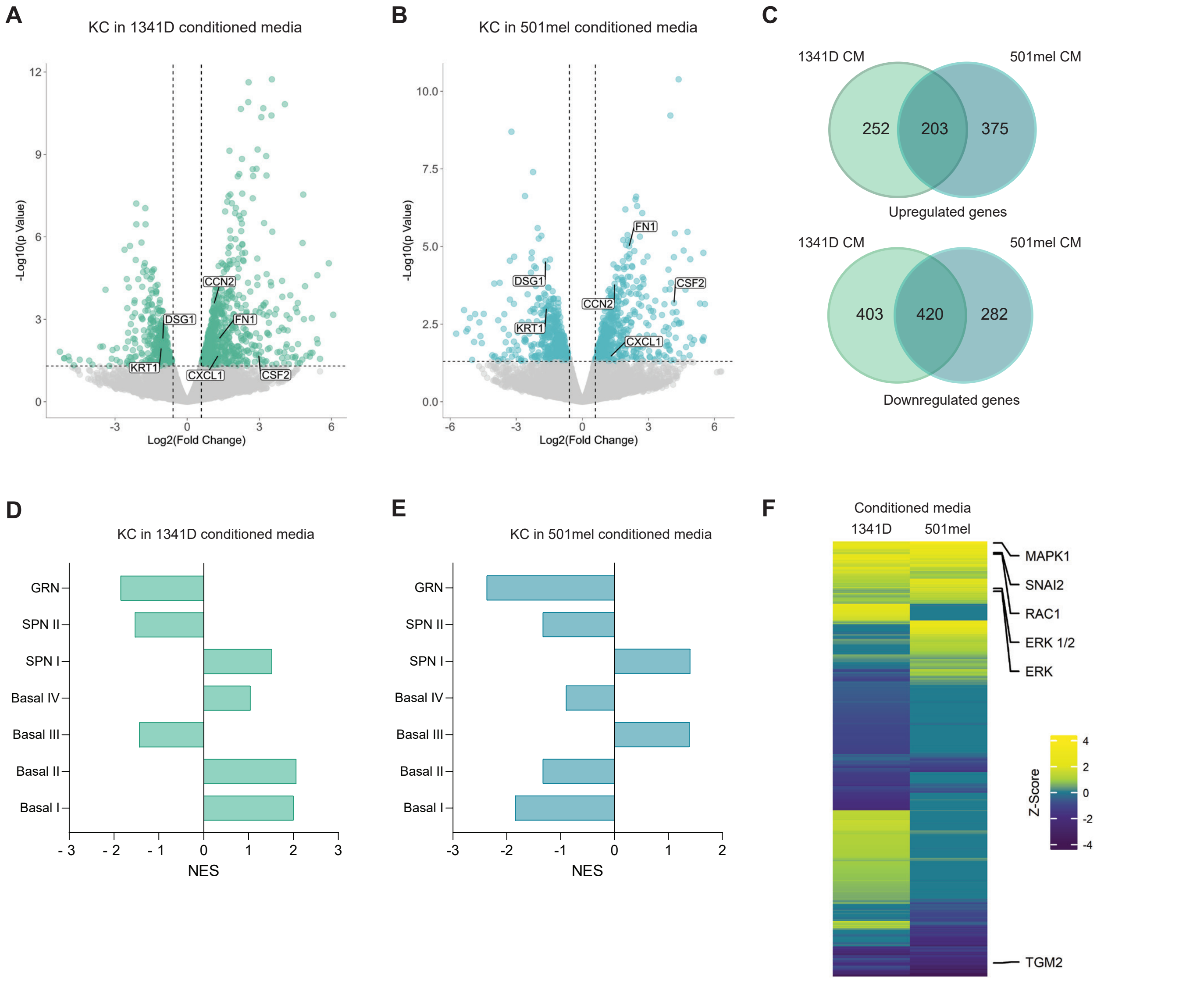


**Supplementary Figure 2. Transcriptomic analysis reveals increased activation of keratinocyte Slug and reduced keratinocyte differentiation signaling by melanoma cells.** A, B) mRNA sequencing was performed on primary human keratinocytes (KC) treated with conditioned media from melanocytes or melanoma cells. Volcano plots depict significantly changes genes in keratinocytes treated with melanoma conditioned media compared to those treated with melanocyte conditioned media. Highlighted genes are those in pathways of differentiation and the identified upstream regulators. C) Venn diagrams depicting overlap between changed genes in those treated with 1341D and 501mel cell lines. D, E) Gene set enrichment analysis (GSEA) comparing differentially expressed gene sets to published signatures of primary human keratinocyte differentiation expression patterns show a decrease in enrichment of granular (GRN) and spinous II (SPNII) layer genes. F) Heatmap showing Upstream Regulators in keratinocytes predicted to be activated by melanoma conditioned media using Ingenuity pathway analysis (IPA) including RAC1, Snai2 and MAPK family signaling members.

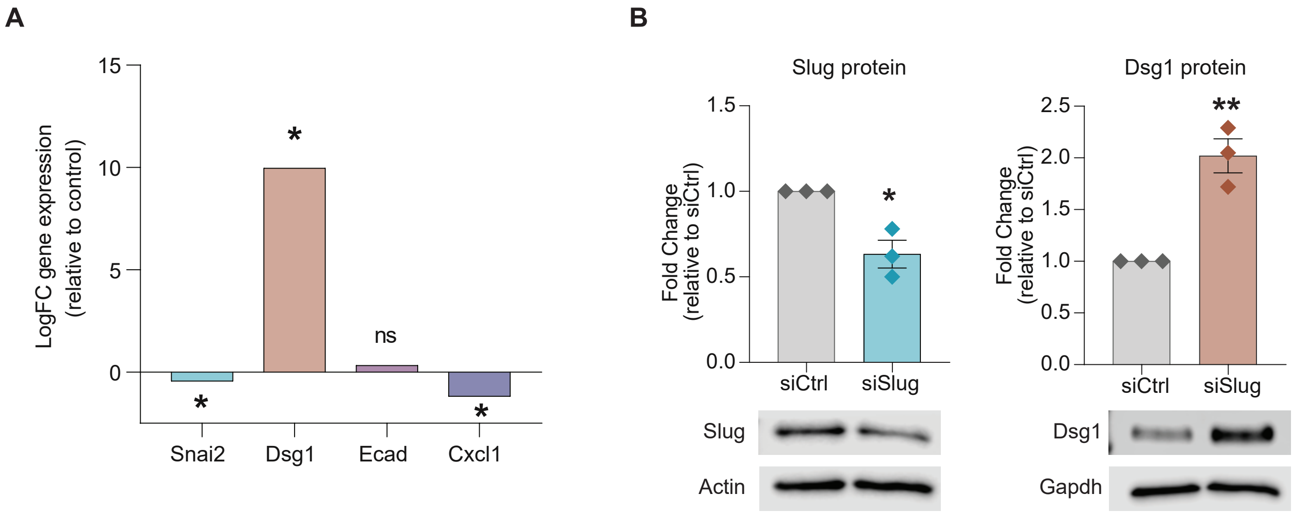


**Supplementary Figure 3. Slug loss in keratinocytes results in decreased Dsg1.** A) Microarray data shows an increase in Dsg1 expression in primary human keratinocytes upon loss of Slug expression, as well as a concurrent decrease in CXCL1 expression. B) siRNA knockdown of Slug causes an increase in Dsg1 protein levels as seen by western blot. n=3 $^*p< 0.05$, $^{**}p< 0.01$. Student's t-test.

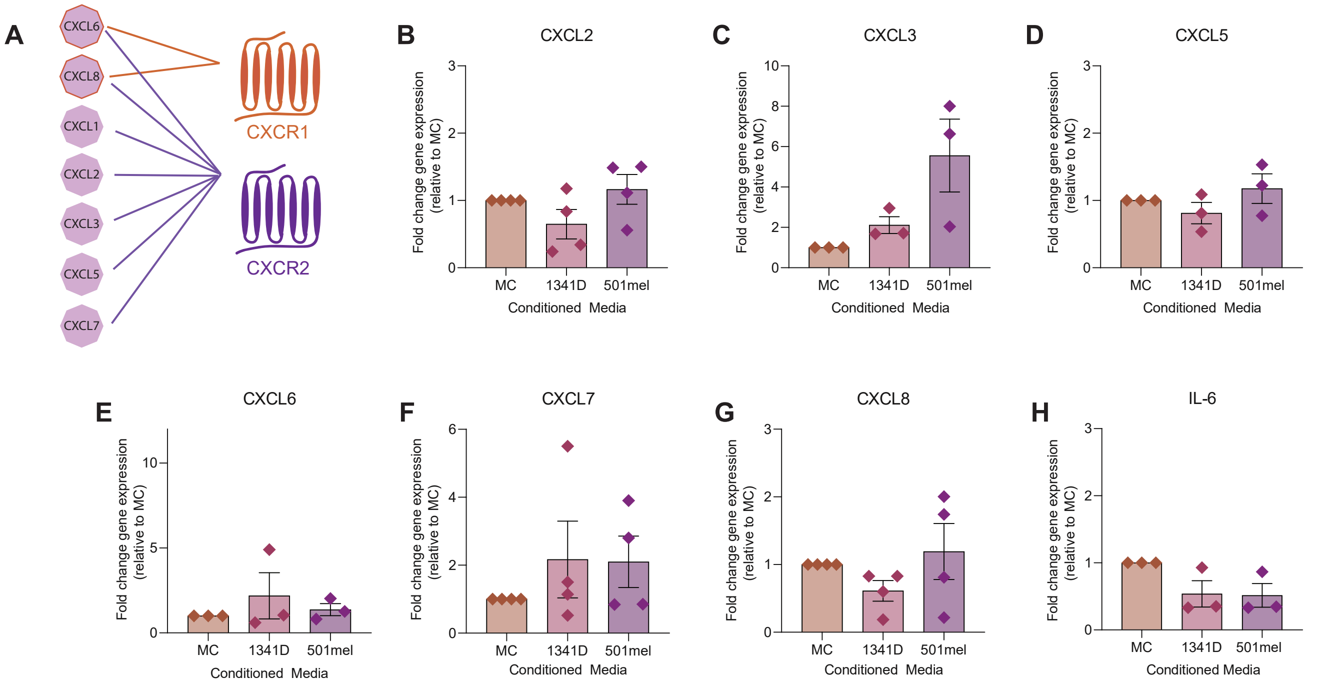


**Supplementary Figure 4. Gene expression of candidate keratinocyte chemokines and cytokines released in response to melanoma conditioned media.** A) Schematic depicting receptor-ligand binding partners for candidate chemokine targets. B-H) Primary human keratinocytes were treated for 48 hours with conditioned media from melanocytes (MC) or the melanoma cell lines 1341D and 501mel and RNA was collected. RT-PCR was performed for chemokines and cytokines induced by shDsg1. Mean +/- SEM depicted. n=3 One way ANOVA.
